# Single trial variability in neural activity during a working memory task: A window into multiple distinct information processing sequences

**DOI:** 10.1101/2022.05.03.490545

**Authors:** Johan Nakuci, Thomas J. Covey, Janet L. Shucard, David W. Shucard, Sarah F. Muldoon

**Author notes:** Johan Nakuci, Sarah F. Muldoon. **Email:**. **Author Contributions:** J.N. and S.F.M designed the analysis; J.N. and T.J.C processed data; T.J.C, J.L.S. and D.W.S. designed the study and acquired data; J.N., T.J.C, J.L.S., D.W.S and S.F.M interpreted the data and wrote the paper. **Competing Interest Statement:** The authors have no competing interests.

## Abstract

Successful encoding, maintenance, and retrieval of information stored in working memory requires persistent coordination of activity among multiple brain regions. It is generally assumed that the pattern of such coordinated activity remains consistent for a given task. Thus, to separate this task-relevant signal from noise, multiple trials of the same task are completed, and the neural response is averaged across trials to generate an event-related potential (ERP). However, from trial to trial, the neuronal activity recorded with electroencephalogram (EEG) is actually spatially and temporally diverse, conflicting with the assumption of a single pattern of activity for a given task. Here, we show that variability in neuronal activity among single time-locked trials arises from the presence of multiple forms of stimulus dependent synchronized activity (i.e., distinct ERPs). We develop a data-driven classification method based on community detection to identify three discrete spatio-temporal clusters, or subtypes, of trials with different patterns of activation that are further associated with differences in decision-making processes. These results demonstrate that differences in the patterns of neural activity during working memory tasks represent fluctuations in the engagement of distinct brain networks and cognitive processes, suggesting that the brain can choose from multiple mechanisms to perform a given task.

**Significance Statement:** Working memory is a complex cognitive ability requiring coordinated activity among multiple brain regions to encode, maintain, and retrieve information. It is generally assumed that the pattern of coordination among brain regions remains consistent and one can average data across multiple trials of the same task. We instead show that there is significant variability in the patterns of brain activity among trials of the same task and develop a method to classify brain activity into distinct subtypes of responses, each with a different spatial and temporal pattern. The subtypes are associated with differences in decision-making processes, suggesting that the brain can use multiple mechanisms to perform a given task.

## Introduction

Variability is a ubiquitous aspect of real-world, day-to-day activities, cognitive testing, and behavioral performance (1–7). Moment-to-moment neurophysiological variability is present during simple and cognitively demanding tasks and may impact the efficiency and accuracy in performance (8–11). However, despite the fact that the presence of variability in neuroimaging data has been accepted, it is still often treated as noise during analysis, as opposed to a feature and richness of the data that could lead to insight into brain function.

Working memory tasks provide an arena to study how neurophysiological variability can relate to cognition. Working memory serves as the scaffolding in which we hold and manipulate thoughts and plan actions. As such, it is a cognitively demanding process requiring coordinated activity among multiple brain regions to encode, maintain, and retrieve information (12, 13). Despite the need to maintain this coordinated activity, variability in neuronal activity across trials is pervasive during a working memory task (6, 14). Such fluctuations are typically considered noise and eliminated through averaging neuronal activity across trials to identify stimulus dependent synchronized activity (15–18).

In electroencephalogram (EEG) recordings, this stimulus dependent synchronized activity can be identified with event-related potentials (ERPs) which provide the average time-locked response to a stimulus locked trial (19). However, averaging trials may obscure important information (19) since trial-to-trial frequency and amplitude differences can arise from (a) fluctuations in postsynaptic response (20), (b) alterations in the balance between excitatory and inhibitory neural systems (21), (c) competition between brain networks (22), and (d) changes in attention (23) as discussed further by Dinstein *et. al* (2015) (24). Additionally, averaging might eliminate the ability to observe distinct forms of synchronized activity (i.e., the ERP) that are engaged differentially across individual trials or contexts, but that are, nonetheless, relevant to behavioral performance.

This issue may be particularly true for tasks with distinct types of trials, or when cognitive processing engages multiple subprocesses that rely on complex interactions between distinct neural networks (25). Therefore, different types of synchronized activity could indicate changes in complex interactions or operations among distributed neural networks during working memory (26).

In the present study, we investigate variability in neural activity across individual trials to determine whether the variability is indicative of distinct forms of stimulus dependent synchronized activity important for working memory and behavioral performance. We use an *n-*back working memory task with three levels of difficulty, 0-back (N_trials_ = 3150, N_subj_ = 21), 1-back (N_trials_ = 3108, N_subj_ = 21), and 2-back (N_trials_ = 3066, N_subj_ = 21), to characterize trial-to-trial variability in neural activity (Fig. 1A). We develop a data-driven clustering method to classify individual trials based on differences in spatial and temporal EEG activity recorded from the scalp. This approach leverages the spatial-temporal resolution of dense-electrode EEG recording to identify distinct characteristics of neuronal activity.

**Figure 1.**
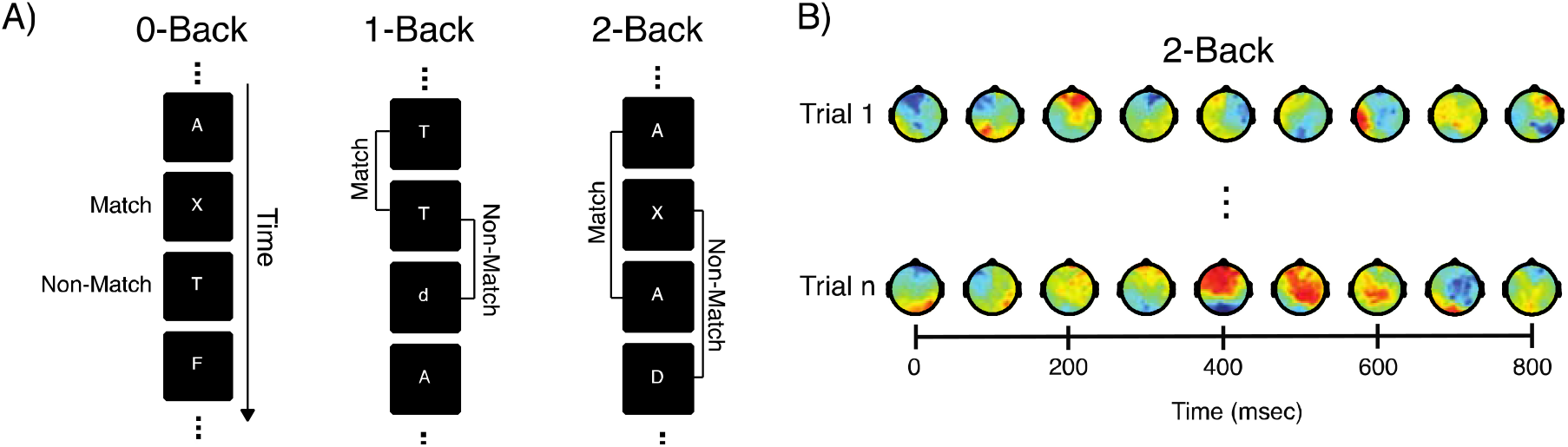
Task description and trial-to-trial variability in neuronal activity. A) An example of the visual 0-, 1- and 2-back task. For the 0-back condition, a match trial was one in which the letter presented was an “X” and the non-match for all other letters. For the 1-back condition, a match trial is one in which the second letter “T” matches the letter one trial before. Similarly, a non-match trial is one in which the letter “C” does not match the letter presented one trial back, “T”. In the 2-back condition, a match trial is one in which the second letter “A” matches the letter presented two trials ago. Similarly, a non-match trial is one in which the letter “D” does not match the letter presented two trials back, “X”. B) EEG activity from two trials during the 2-back condition from stimulus onset (0 msec) to 800 msec from the same subject. The brain activity between the trials exhibits stark differences particularly after 200 msec.

## Results

During the *n*-back working memory task, trial-to-trial variability is marked by the presence of different spatial and temporal patterning in neuronal activity (Fig. 1B). We investigated if these differences in neural activity indicated the existence of distinct forms of stimulus dependent synchronized activity. We developed a data-driven clustering method based on community detection that utilizes topographical and temporal differences to classify individual trials. The pairwise similarity in neuronal activity across trials was determined using the Pearson Product Correlation of full brain EEG sensor activity from stimulus onset (0 msec) to 852 msec (144 sensors × 252 data points per sensor) regardless of trial type. This trial-by-trial similarity matrix was then clustered using Modularity-Maximization, an unsupervised clustering method that does not require the number of clusters to be specified (27). As shown in Fig. 2A for the 2-back task, modularity-maximization separated individual trials into 3 clusters, or subtypes we label T1 (N_trials_ = 1134), T2 (N_trials_ = 1061), and T3 (N_trials_ = 871)(Fig. 2A).

**Figure 2.**
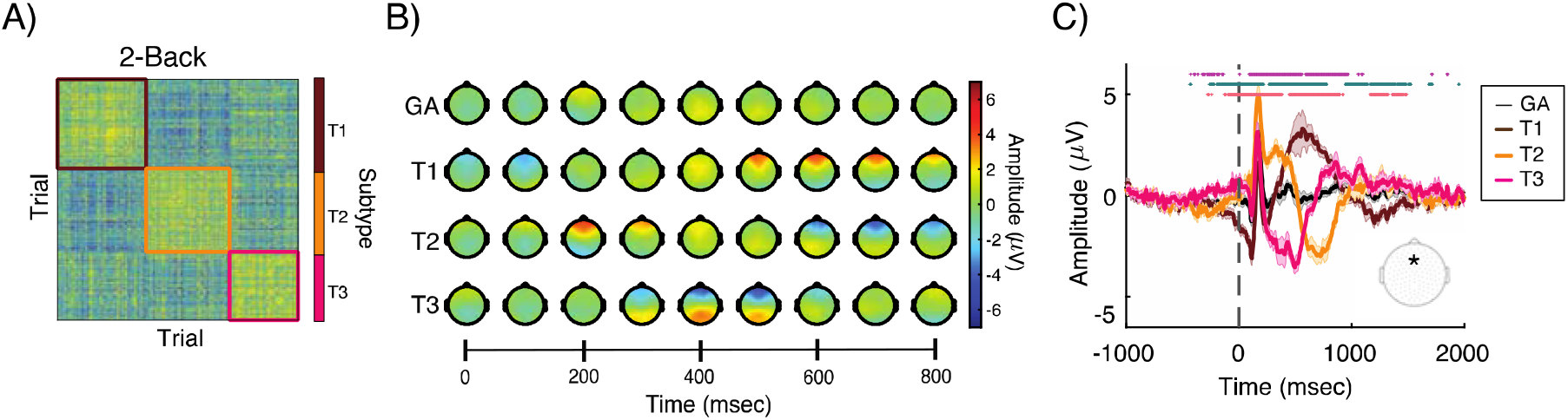
Subtypes of individual trials in the 2-back. A) Modularity-Maximization based clustering identified 3 subtypes of trials, T1, T2, and T3. The colored squares correspond to the trials composing each subtype. Pearson Correlation was used to calculate the spatial temporal similarity of the EEG activity from individual trials. B) Spatial and temporal ERP activity from the T1, T2, T3 subtypes and the grand average (GA) at 9 time points post-stimulus, respectively. C) ERP activity from frontal-midline electrode. Each waveform shows the mean (thick line) and standard error of the mean (shaded area). Insert shows the location of the EEG electrode from which the ERP waveforms were derived. Statistical testing is based on Paired T-test and FDR corrected for multiple comparison (*p_corrected_ < 0.05). Statistically significant differences in amplitude are marked at the top of each panel (the three rows of colored asterisks) between: * T1 and T2; * T1 and T3; * T2 and T3.

ERPs are neuronal activity derived from spontaneous and synchronized activity generated in response to a stimulus and averaged across trials for a time-locked task (19). To better assess the potential novelty of the analytical approach used here, we examined the correspondence between the spatial and temporal properties of the conventional signal-averaged ERP waveform (the grand average (GA) generated by averaging across all trials regardless of trial type) and the ERPs acquired by separately averaging across trials within each subtype (T1, T2, T3) obtained from the modularity-maximization analysis (Fig. 2B-C). Qualitatively, the topography of the ERPs from each subtype in the 2-back condition fluctuated between frontal and parietal positive amplitude relative to each other and the grand average (Fig. 2B; SI Movie 1). Further, these topographical fluctuations show significantly different temporal profiles among each other when focusing on a single EEG electrode (Fig. 2C).

Expanding this analysis to the 0- and 1-back condition, similarly, modularity-maximization identified 3 subtypes for the 0-back [T1 (N_trials_ = 951), T2 (N_trials_ = 1101), and T3 (N_trials_ = 1098)] and 1-back [T1 (N_trials_ = 899), T2 (N_trials_ = 987), and T3 (N_trials_ = 1222)] (Fig. 3A and D, respectively). The topography of these ERPs fluctuated between frontal and parietal positivity (Fig. 3B and E, SI Movie 2 & 3, respectively) and with significant amplitude differences from a single electrode, reflecting the findings from the 2-back (Fig. 3C and F, respectively). Notably, each of these distinct waveforms was observed for different trials within a given individual indicating that the subtypes are not simply due to differences between individuals (SI Fig. 1).

**Figure 3.**
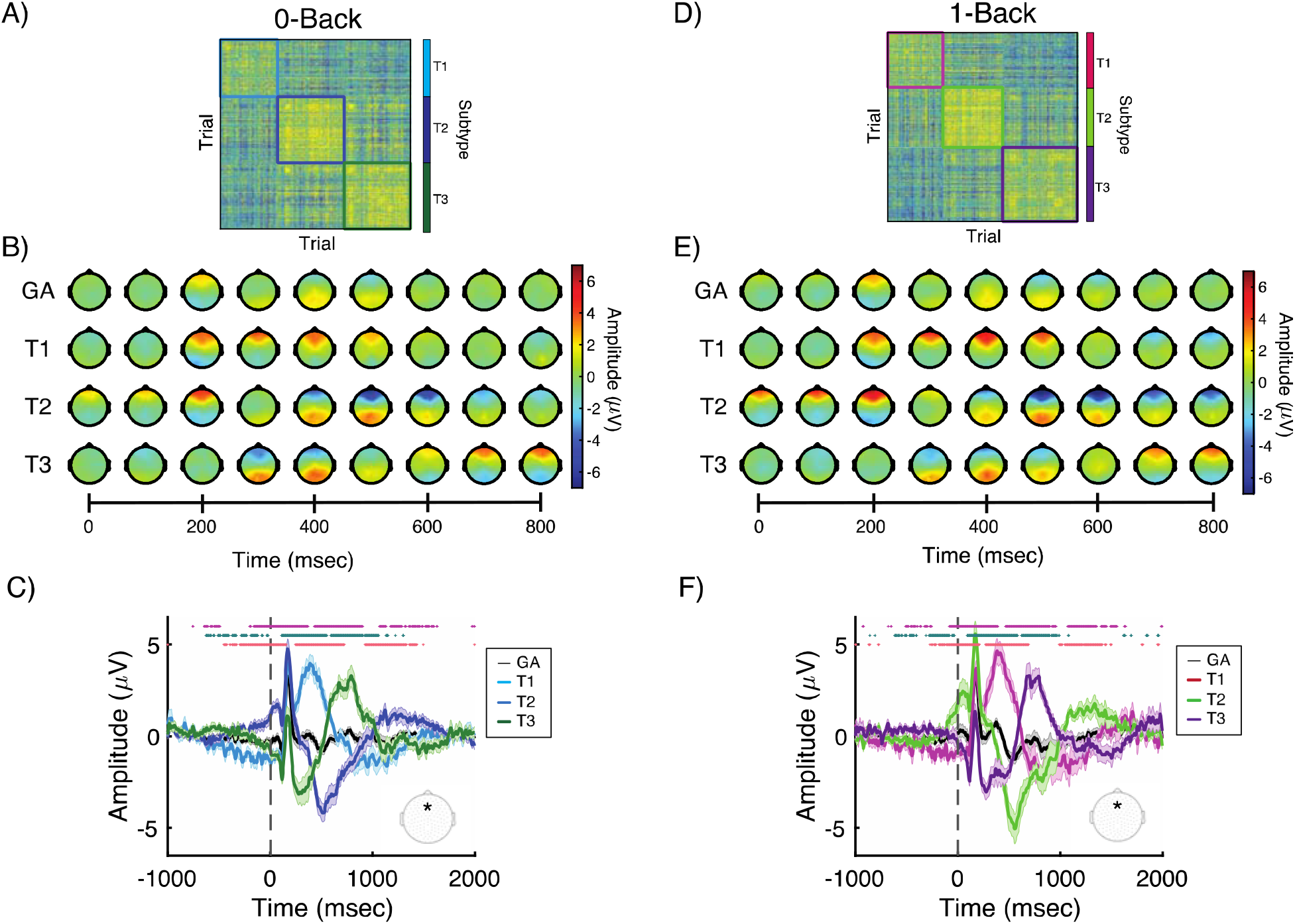
Subtypes of individual trials in the 0 and 1-back. A and D) Modularity-Maximization based clustering identified 3 subtypes of trials, T1, T2, and T3 in 0- and 1-back, respectively. The colored squares correspond to the trials composing each subtype. B and E) Spatial and temporal ERP activity from the T1, T2, T3 subtypes and the grand average (GA) at 9 time points post-stimulus, respectively. C and F) ERP activity from frontal-midline electrode. Each waveform shows the mean (thick line) and standard error of the mean (shaded area). Insert shows the location of the EEG electrode from which the ERP waveforms were derived. Statistical testing is based on Paired t-tests and FDR corrected for multiple comparison (*p_corrected_ < 0.05). Statistically significant differences in amplitude are marked at the top of each panel (the three rows of colored asterisks) between: * T1 and T2; * T1 and T3; * T2 and T3.

We quantified the spatial-temporal similarity between ERPs from subtype to traditional grand-average to determine if subtypes contain ERPs different from the average. The spatial-temporal similarity was estimated using the Pearson Correlation from 0msec to 852msec. The grand-average ERP was strongly (r > 0.75) correlated with only one of the ERP subtypes in the 0-back (SI Fig. 2A, *Middle*) and 1-back (SI Fig. 2B, *Middle*), and moderately (r > 0.50) correlated with one subtype in the 2-back (SI Fig. 2C, *Bottom*).

The above analysis provides only an overarching similarity between the grand averages and the subtypes. We therefore focused on identifying the specific time points in which the ERPs differ by quantifying the similarity at each time point between subtypes and the grand average. For all ERP subtypes, there was a period of high topographical similarity (r > 0.75) between approximately 152msec to 200msec (Fig. 4A-C). This period is associated with visual processing and attention indicating that the divergence among the subtype waveforms is not due to differences in early visual processing or differences in attention at stimulus onset (29).

**Figure 4.**
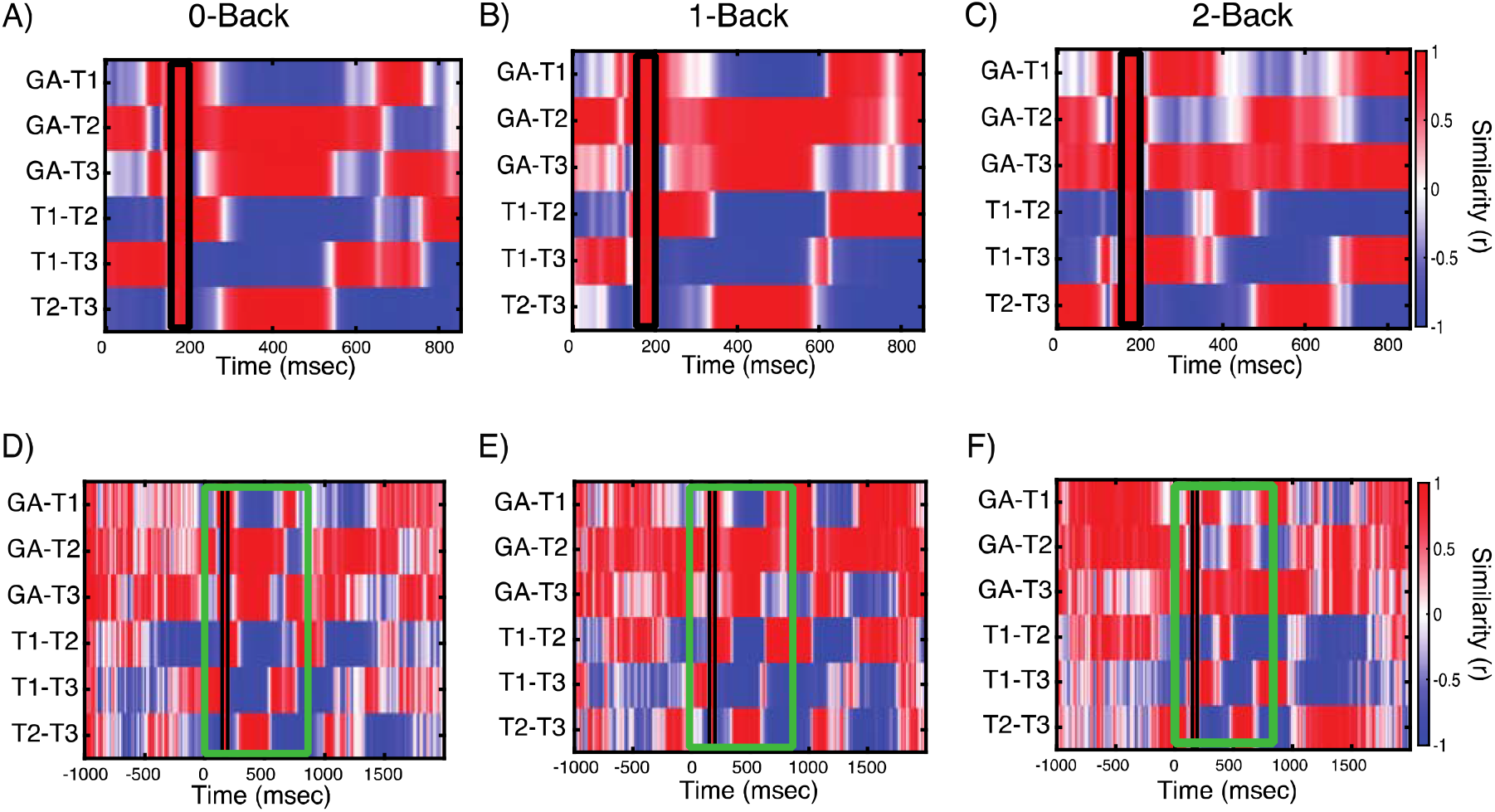
Spatial and temporal heterogeneity in ERP active from each of the subtypes. A-C) The similarity in spatial topography using the Pearson Product Correlation among the ERP subtypes and the grand average for the 0-, 1- and 2-back, respectively. The spatial similarity flips between positively and negatively correlated epochs with a brief period between with high spatial similarity among the GA and subtypes (black box). The spatial similarity was calculated at each time point between stimulus onset (0msec) to 852 msec. D-F) The spatial similarity from -1000 to 2000msec post-stimulus. In panels D-F, the green box corresponds to the period in panel A-C.

Outside of this approximately 50msec window, the ERP waveforms fluctuated between positively and negatively correlated topographies suggesting the different ERP waveforms are the result of brain regions/networks becoming active at different periods in the trial. These topographical fluctuations extended to the pre-stimulus and after 852 msec, indicating they were not an effect of the time range used for classification (Fig. 4D-F). The presence of these fluctuations in the pre-stimulus time range suggests that pre-stimulus activity might play a role in determining post-stimulus activity (30) and behavioral performance (31–34).

The existence of these subtypes raised the possibility that they were relevant for behavioral performance. We tested for differences in reaction times and the proportion of correct responses for match/non-match trials among subtypes. We found significant differences in reaction times among subtypes in the 0-back (T1 = 558 ± 209.75; T2 = 513 ± 219; T3 = 506 ± 209.25 [Median ± IQR msec]; Kruskal-Wallis Test, χ^*2*^ = 43.56, p = 3.47 × 10^−10^), 1-back (T1 = 699 ± 261.75; T2 = 624 ± 257.50; T3 = 620 ± 302 [Median ± IQR msec]; Kruskal-Wallis Test, χ^*2*^ = 92.51, p = 8.14 × 10^−21^), and 2-back (T1 = 767.50 ± 374; T2 = 747 ± 325.25; T3 = 705 ± 368.5 [Median ± IQR msec]; Kruskal-Wallis Test, χ^*2*^ = 18.8, p = 7.95 × 10^−5^) suggesting subtypes were associated with decision making processes (Fig. 5A-C). However, these processes were not associated with stimulus categorization because the subtypes contained the same proportion of correct match/non-match trials (ANOVA, p > 0.05; Fig. 5D-F). Thus, not only do the subtypes appear to have ERPs with spatial-temporal activity that differ from each other and from the traditional grand-average, they also are associated with differences in behavioral performance. This finding supports work suggesting the traditional grand average ERP, in the best case, captures only a portion of true stimulus-driven synchronized brain activity and in the worst case, the grand average ERP obscures the actual brain activity associated with cognitive processing (19, 28).

**Figure 5.**
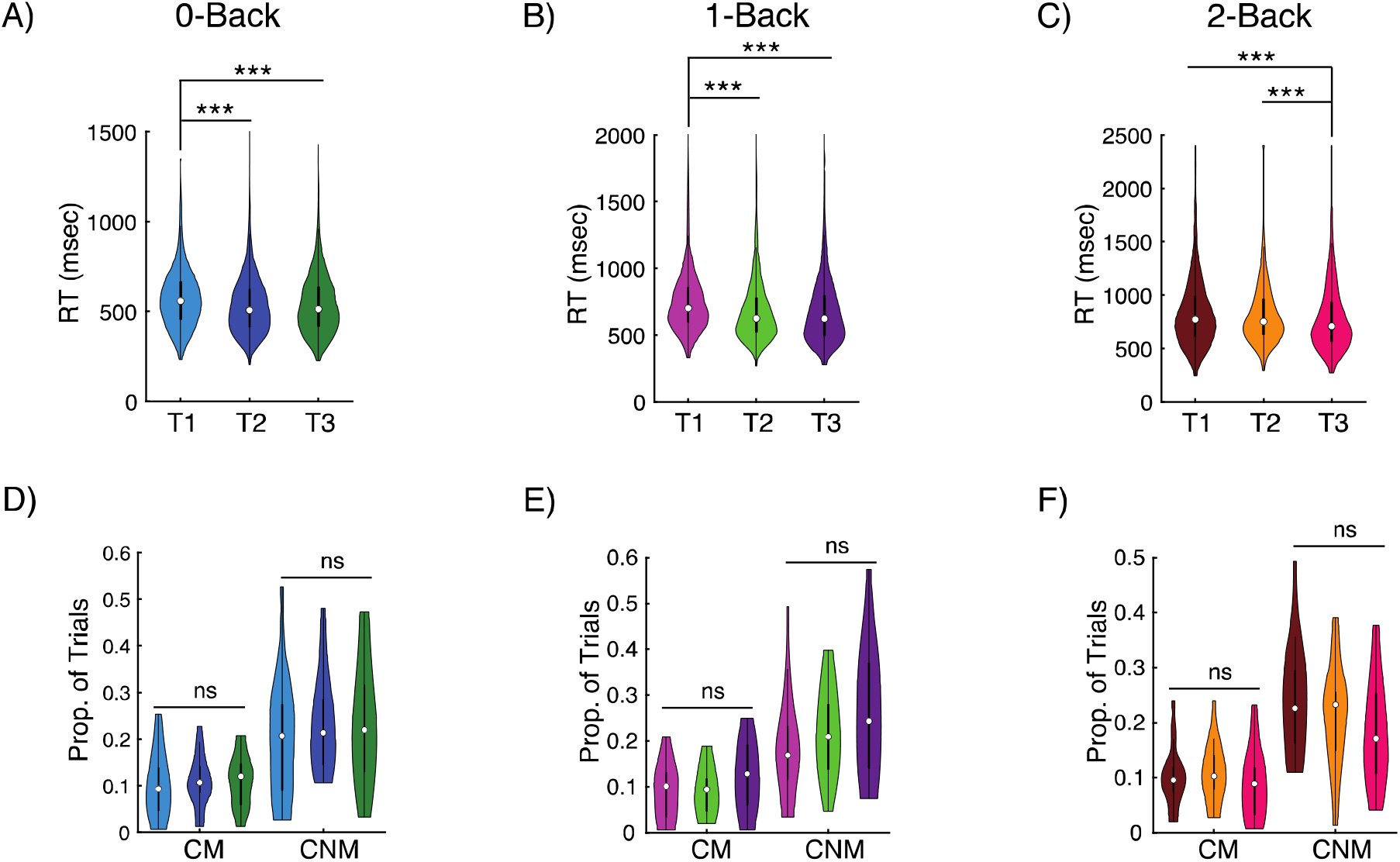
Behavioral associations among trial subtypes. A-C) Differences in reaction times between trial subtypes for the 0-, 1-, 2-back, respectively. Post-hoc statistical tests were performed with Wilcoxon rank sum test and Bonferroni multiple comparison correction. D-F) The proportion of correct match (CM) and correct non-match (CNM) responses was equivalent between trial subtypes for the 0-, 1-; and 2-back, respectively. Violin plot shows the distribution of the data with the circle representing the median value, box containing the 25^th^ and 75^th^ percentiles and line representing all data that are not considered outliers. *** p_corrected_ < 0.001, ns = not significant.

We next investigated if the spatial temporal activity of the ERP subtypes were similar across the *n*-back conditions. In each condition, subgroups are labeled T1, T2, and T3 in decreasing order of the reaction time associated with the subtype (i.e., T1 is always the subtype associated with the slowest reaction time). As seen in Fig. 6A, when comparing the spatial-temporal profiles of each subtype with the corresponding subtype (i.e., T1 vs T1, T2 vs. T2, T3 vs. T3), we found strong (r > 0.75) similarity in spatio-temporal patterning of subtypes between 0- and 1-back tasks. When comparing the 2-back task with both the 0- and 1-back tasks (Fig. 6B-C), we still observe moderate (r > 0.50) similarity between corresponding T2 and T3 subtypes, although the similarity for T1 subtypes weakens significantly. Interestingly, we observe the lowest values of similarity when comparing 0- and 2-back tasks, suggesting that the difficulty of the task modifies the spatio-temporal profile of the subtyped ERPs.

**Figure 6.**
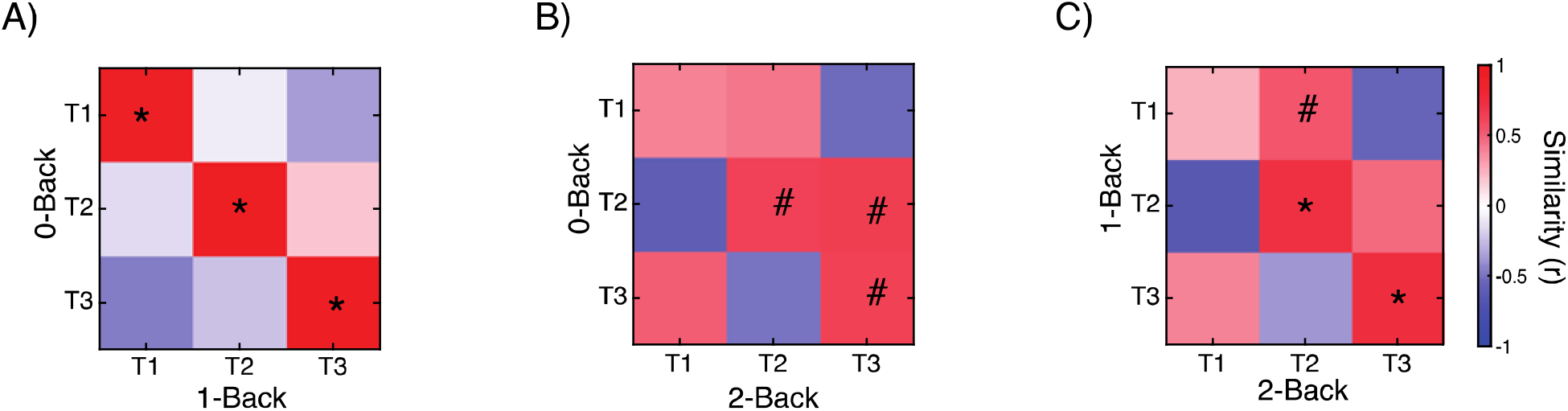
Similarity of ERP subtypes among *n*-back conditions. Spatial-temporal ERP similarity among subtypes between the A) 0- and 1-back condition; B) 0- and 2-back condition; and C) 1- and 2-back condition. The spatial-temporal similarity was estimated using Pearson Correlation from 0msec to 852msec. * r > 0.75; # r > 0.50. p < 10^−15^

## Discussion

Trial-to-trial variability is ubiquitous in neurophysiological activity during simple and cognitively demanding tasks (35, 36). This situation raises questions about the role neuronal variability plays in brain function and cognition. Variability in neuronal activity across different stimulus categories may provide a benefit in that it allows flexibility when responding to changing cognitive demands (37). However, this variability may need to be calibrated appropriately to produce a beneficial response because otherwise it could be detrimental to neural performance (38–40) and/or may reflect underlying functional/structural disturbances (24, 41–43). In the present study, we investigated if variability in neural activity indicates the existence of distinct forms of stimulus dependent synchronized activity among trials reflecting different cognitive processes in healthy individuals.

We developed a data-driven topographical method to classify neuronal activity measured with EEG from individual trials during a visual *n*-back task, with three levels of difficulty or working memory load: 0-, 1-and 2-back. This analysis allowed us to define not only a standard GA ERP, but also ERPs for each of the subtypes identified from the clustering analysis, in order to investigate the relation between stimulus driven synchronized activity and working memory and behavioral performance. We limited our analysis primarily to ERPs because the ERP is a well characterized form of synchronized brain activity generated in response to a stimulus (19).

We found that trial-to-trial variability in neuronal activity was limited to a few distinct waveforms or subtypes and these subtypes contained ERPs with unique spatial and temporal organization. Further, the distinct subtypes were associated with differences in decision-making processes. It is often assumed that multiple successive components of the ERP represent successive steps of information processing, from early sensory/perceptual discrimination through target categorization and memory maintenance functions. However, a major concern and limitation in ERP research is that multiple components may be superimposed over each other. Thus, components may not always occur together on the same trial, but rather occur on different trials, which then, when averaged, appear as though they are occurring in temporal succession. Our results indicate that such concerns are well founded.

Spatial and temporal differences among subtype ERP waveforms were not due to differences in the early visual processing and attention since the period between 152-200msec was marked by high topographical similarity. This period of similarity in neuronal activity is in line with previous work using linear discriminant analysis to classify individual-trials in which the early ERP component was not associated with a participant’s ability to discriminate between choices (44, 45). Therefore, the divergence among the subtypes/ERP waveforms was not due to differences in the early visual processes or differences in early attention, but likely due later stage cognitive processes.

Outside of this 50 msec period, the neurophysiological activity fluctuated between positively- and negatively correlated ERP topographies. For example, the topographies of the T1 and T2 subtypes in the 2-back were negatively correlated from approximately 200 to 300 msec and positively correlated between 400 to 500 msec post-stimulus. This “flickering” could be a marker of distinct brain networks and cognitive processes being recruited to accomplish the task. Engagement of different cognitive processes and their corresponding brain regions could be a result of having to make a decision under varying levels of uncertainty or conflict between choices (46). In the *n*-back task, uncertainty and conflict could arise from the maintenance and/or decision component because, unlike many other working memory tasks, the *n*-back integrates these two components simultaneously (47).

Furthermore, these unique forms of stimulus-driven brain activity (subtypes) were associated with behavioral performance. Trials with slowed response speed were associated with ERP subtypes that contained a positive frontal component at approximately 300 msec, which is aligned with the canonical spatial-temporal characteristics of the novelty-P3 (48) and with enhancement of executive function associated with increased task difficulty (49). The novelty-P3 is associated with equivocation during cognitive performance suggesting participants may be perceiving a stimulus that they did not expect (48, 50). On the other hand, efficient (faster) performance was associated with frontal negativity at ∼250 msec, the N2-P3 complex (51), and followed by frontal positivity at ∼400 msec, the N400-late positivity complex (52). The combination of neurophysiological and behavioral effects supports the conclusion that distinct brain networks and cognitive processes are recruited to accomplish the task.

Interestingly, and perhaps not too surprisingly, we did observe a strong similarity in the spatio-temporal profiles of ERP subtypes between the 0- and 1-back tasks; both are tasks with low difficulty. While further work should investigate the exact cognitive processes involved in each subtype, this finding suggests that the brain switches between similar cognitive processes for each of these tasks. The decrease in similarity when comparing subtasks from the 0- or 1-back task with the 2-back task suggests that task difficulty might also modify the exact cognitive processes involved. It is also interesting to observe that, the spatio-temporal profile of the T2 and T3 subtypes are more similar between the 0- and 2-back tasks, whereas the T1 and T2 subtypes show increased similarity between the 1- and 2-back tasks. Future work should investigate more specifically how task difficulty modifies these spatio-temporal profiles and how these profiles relate to the cognitive processes involved in performing the task.

Lastly, although the analysis was limited to EEG and ERPs, our methodological and analytical approach can be applied to other high-dimensional neurophysiological recordings such as individual neurons (53). Additionally, this analytical approach also can be applied to activity from frequency bands obtained during task-induced EEG activity, another widely used electrophysiological method for evaluating cognitive function (54).

In summary, these results indicate that working memory is dependent on a set of limited, but diverse sequences of cognitive processes reflecting coordinated neuronal activity from spatially and temporally diverse regions. Why the brain might benefit from having multiple but distinct processes to perform the same function remains an open question. One might speculate that it could relate to robustness of task performance or depleted metabolic resources for a given brain region over time. Alternatively, the multiple processes might arise from how efficiently the brain processes the stimulus. For example, if there is noise or uncertainty in the processing of the stimulus, the brain might engage in a different set of processes than if the uncertainty is not present. Regardless, our results indicate that in order to answer these questions, it will be necessary to perform a more nuanced analysis of neuroimaging data during task performance, as valuable information is lost by averaging across trials. This loss can be mitigated through classifying individual trials to estimate the “real” stimulus-driven synchronized neuronal activity contributing to working memory and cognitive function, and we encourage future work to embrace this approach.

## Materials and Methods

### Participants and task

A total of 41 healthy participants completed a visual *n-back* task. Participants were recruited as part of a larger study examining cognitive disturbances related to Systemic Lupus Erythematosus. The current analyses were based on 21participants (17 Female; Age = 42 ± 12 years) who completed a visual *n-back* task (0-, 1-, 2-back). The number of participants in the current analysis was lower than that of the total number in the study because participants were removed if more than 5% of the trials from any of the n-back tasks exceeded 100 *μ*V which likely was due to artifacts such as eye-blinks and movement. The removal of participants instead of trials, ensured that all participants had the same number of trials for analysis and contributed equally to the clustering analysis. All participants were required to have at least 50 percent correct trials to be included in the study.

All participants analyzed in the present study were neurologically normal. Participants were excluded if they had history of head trauma, visual problems that could not be corrected, learning disability, neurological impairment, or if they had an Axis I psychiatric disorder. Participants provided informed consent and received financial compensation for participating in the study. This research was approved by the University at Buffalo Institutional Review Board.

Each participant completed a visual n-back task (0-, 1-, 2-back) with a stimulus duration of 400ms, and 2000ms inter-stimulus-interval. Letters were presented at the center of a computer monitor, approximately 70 cm from the participant’s nasion. The *n-back* conditions were administered in the following order: 0-back, 1-back, and 2-back. A practice trial preceded each condition. In each of the conditions a total of 150 letters were presented one at a time on a computer screen via E-Prime software (E-Prime, 2002). Each condition was divided into two blocks of 75 letters. A brief break was taken between blocks and an approximately 5-minute break was taken between conditions. During the task, participants were instructed to press the inner buttons of a four-button response pad, simultaneously with both thumbs, if the letter they observed on the screen matched the letter presented *n* trials back. They pressed the outer buttons of the response pad with both thumbs if the letter presented on the screen did not match the letter presented *n* trials back. For the 0-back condition, participants were instructed to press the “match” buttons if an “X” was presented, and the “non-match” buttons for all other letters. Fifty of the letters in each condition were match trials and 100, 98, and 96 were a non-match trials in the 0-, 1-, and 2-Back conditions, respectively (the first letter of each block for the 1-back, and the first two letters of each block for the 2-back were not counted due to the task requirements). Reaction time (RT) to both match and nonmatch trials, correct responses, and EEG were simultaneously recorded for each condition.

### EEG recording and processing

Participants were seated in a comfortable chair in a dimly lit, sound attenuated room in front of a computer monitor. A 256-channel dense electrode HydroCel Geodesic Sensor Net (Phillips-Electrical Geodesics, Inc., Eugene, OR) was fitted onto the participant’s head. During EEG acquisition, electrode impedances were kept below 50 kΩ whenever possible. A vertex reference was used during recording, and EEG data were filtered between 0.1 to 100 Hz and digitized at a sampling rate of 250 Hz.

The EEG cleaning procedure was implemented using EEGLAB v14 (55). EEG data were band-pass filtered between 0.5 Hz and 50 Hz using a Finite Impulse Response filter. Next, artifact subspace reconstruction (ASR) was used to determine and remove bad channels (56). Following ASR, independent component analysis (ICA) decomposition, and ADJUST was used to remove ICA components associated with noise (eye blinks, muscle artefact, etc.) (57). Following ICA decomposition, bad channels were interpolated using spherical interpolation.

ASR was used to interpolated data segments not cleaned by ICA. Data was scanned with a 500 msec sliding window to find segments that are greater than four standard deviations from the reference segment. These noisy segments are treated as missing data and reconstructed. Lastly, the data were re-referenced to a common average reference and were segmented from -1000 msec to 2000 msec post-stimulus, and baseline corrected by subtracting the mean amplitude of the baseline portion (−1000 to 0 msec) of the epoch.

### Modularity-maximization clustering analysis

Prior to clustering, the first two trials in the 2-back were removed because they were non-response trials. Next, data were pooled from all 21 participants resulting in 3066 trials for the 2-back. A trial-by-trial similarity matrix was created using the Pearson Product Correlation between trials calculated from 0 msec (stimulus onset) to 852 msec across 144 sensors above the canthomeatal line, corresponding to 36,288 data points per trial. Clustering was limited to an interval from 0 msec to 852 msec post-stimulus because: 1) this time interval contains stimulus driven activity; 2) the 852 msec time occurs after the median reaction time of 745 msec, ensuring the relevant brain activity related to the trial that leads to the eventual behavioral response is maintained; and 3) this minimized interference from EEG artifacts (i.e. eye blinks) that tended to occur most frequently in the inter-stimulus-interval (after 852 msec).

Clustering of the similarity matrix was conducted using modularity-maximization (27). Modularity-maximization identifies clusters in which the similarity within a cluster is greater than between clusters. In addition, modularity-maximization does not require the number of clusters to be specified. The resolution of the cluster can be controlled with resolution parameter; here we use γ = 1. Modularity-maximization was implemented with the Generalized Louvain algorithm part of the Brain Connectivity Toolbox (58). Clustering results from the Generalized Louvain algorithm can be dependent on the initial seeding. To avoid such suboptimal results, clustering was repeated 100x and final clustering results were based on the consensus across iterations (59).

### Identification of event-related potentials (ERPs)

ERPs for each subtype were generated by averaging all trials within each of the corresponding clusters. In addition to ERP Subtypes, grand averaged ERPs were generated to assess the similarity in topography obtained between a standard approach to ERP analysis and that of the clustering methodology outlined above. Grand averaged ERPs were generated from averaging all trials collapsed in the 2-back task.

### Statistical analysis and software

Statistical testing used both parametric and non-parametric tests. To test for ERP amplitude differences, we used Paired t-test and False Discovery Rate (FDR) multiple comparison correction. Statistical differences for reaction times were tested using Kruskal-Wallis test and post-hoc comparison was conducted using Wilcoxon rank sum test with Bonferroni multiple comparison correction because of the non-parametric distribution of the corresponding values. Whereas differences in accuracy were tested with ANOVA. All data processing and statistical analyses were performed in MATLAB 2018b.

## Data Sharing

All data and code is available via request to the authors upon publication.

## Acknowledgments

This work was supported by National Institutes of Health Grant NS049111, PI JLS.

